# Refining pairwise sequence alignments of membrane proteins by the incorporation of anchors

**DOI:** 10.1101/2020.09.16.299453

**Authors:** René Staritzbichler, Edoardo Sarti, Emily Yaklich, Antoniya Aleksandrova, Markus Stamm, Kamil Khafizov, Lucy R Forrest

## Abstract

The alignment of primary sequences is a fundamental step in the analysis of protein structure, function, and evolution. Integral membrane proteins pose a significant challenge for such sequence alignment approaches, because their evolutionary relationships can be very remote, and because a high content of hydrophobic amino acids reduces their complexity. Frequently, biochemical or biophysical data is available that informs the optimum alignment, for example, indicating specific positions that share common functional or structural roles. Currently, if those positions are not correctly aligned by a standard pairwise alignment procedure, the incorporation of such information into the alignment is typically addressed in an ad hoc manner, with manual adjustments. However, such modifications are problematic because they reduce the robustness and reproducibility of the alignment. An alternative approach is the use of restraints, or anchors, to incorporate such position-matching explicitly during alignment. Here we introduce position anchoring in the alignment tool AlignMe as an aid to pairwise sequence alignment of membrane proteins. Applying this approach to realistic scenarios involving distantly-related and low complexity sequences, we illustrate how the addition of even a single anchor can dramatically improve the accuracy of the alignments, while maintaining the reproducibility and rigor of the overall alignment.

## Introduction

Comparison of primary sequences of proteins and nucleic acids is a fundamental step in analyzing evolutionary relationships and conservation of functional properties. Solutions to the problem of primary sequence alignment have typically focused either on identifying optimally-matched short fragments or on obtaining the best agreement between two entire sequences, which are referred to as local and global alignments, respectively. In both cases, the likelihood of matching of each pair of amino-acids in the alignment is scaled according to a reference table of amino-acid substitution rates, such as the BLOSUM or PAM matrices (Henikoff and Henikoff 1992; Dayhoff, Schwartz, and Orcutt 1978), or in the case of α-helical membrane proteins, the JTT, PHAT, or SLIM matrices (Jones, Taylor, and Thornton 1994; Ng, Henikoff, and Henikoff 2000; Müller, Rahmann, and Rehmsmeier 2001). For a global alignment, the cost of introducing gaps of different lengths must also be considered.

From an algorithmic perspective, global alignments are commonly achieved using the Needleman-Wunsch algorithm (Needleman and Wunsch 1970) or hidden Markov Models (Remmert et al. 2012; Eddy 2011). The classical Needleman-Wunsch approach involves first filling out a so-called dynamic programming matrix reflecting the scores of all possible pairs of matching positions and then identifying a high-scoring, optimal path through that matrix. When the overall similarity of the two proteins is high, for example, with > 50% identical residues, achieving accurate global alignments is relatively trivial with current approaches (Rost 1999). At lower levels of similarity, accuracy can be improved by incorporating representations of properties such as secondary structure (Tang et al. 2003; Zhou and Zhou 2005; Pei and Grishin 2007; Dong et al. 2008; Deng and Cheng 2011) or, in the specific case of membrane proteins, the membrane-spanning regions (Pirovano, Feenstra, and Heringa 2008; Stamm et al. 2013; Bhat et al. 2017).

Even with the aforementioned advances, membrane proteins continue to present a significant challenge to pairwise sequence alignment methods, for two reasons. First, the regions that span the hydrophobic core of the lipid bilayer are highly hydrophobic and therefore their composition tends to be low in complexity (Hedman et al. 2002). Second, membrane proteins are thought to be among the most ancient of all proteins (Sojo et al. 2016), and consequently the relationships between membrane proteins – even those with similar structural folds – can be very weak and may also be complicated by the addition of peripheral segments (**Fig. 1**). Consequently, even state-of-the-art methods such as PRALINE™ (Pirovano, Feenstra, and Heringa 2008), HHalign (Söding 2004), TM-Coffee (Chang et al. 2012) and AlignMe (Stamm et al. 2013), fail to correctly align large fractions of the positions in the most dissimilar pairs of sequences in test sets such as BAliBASE2-ref7 (Bahr et al. 2001).

**Figure 1.**
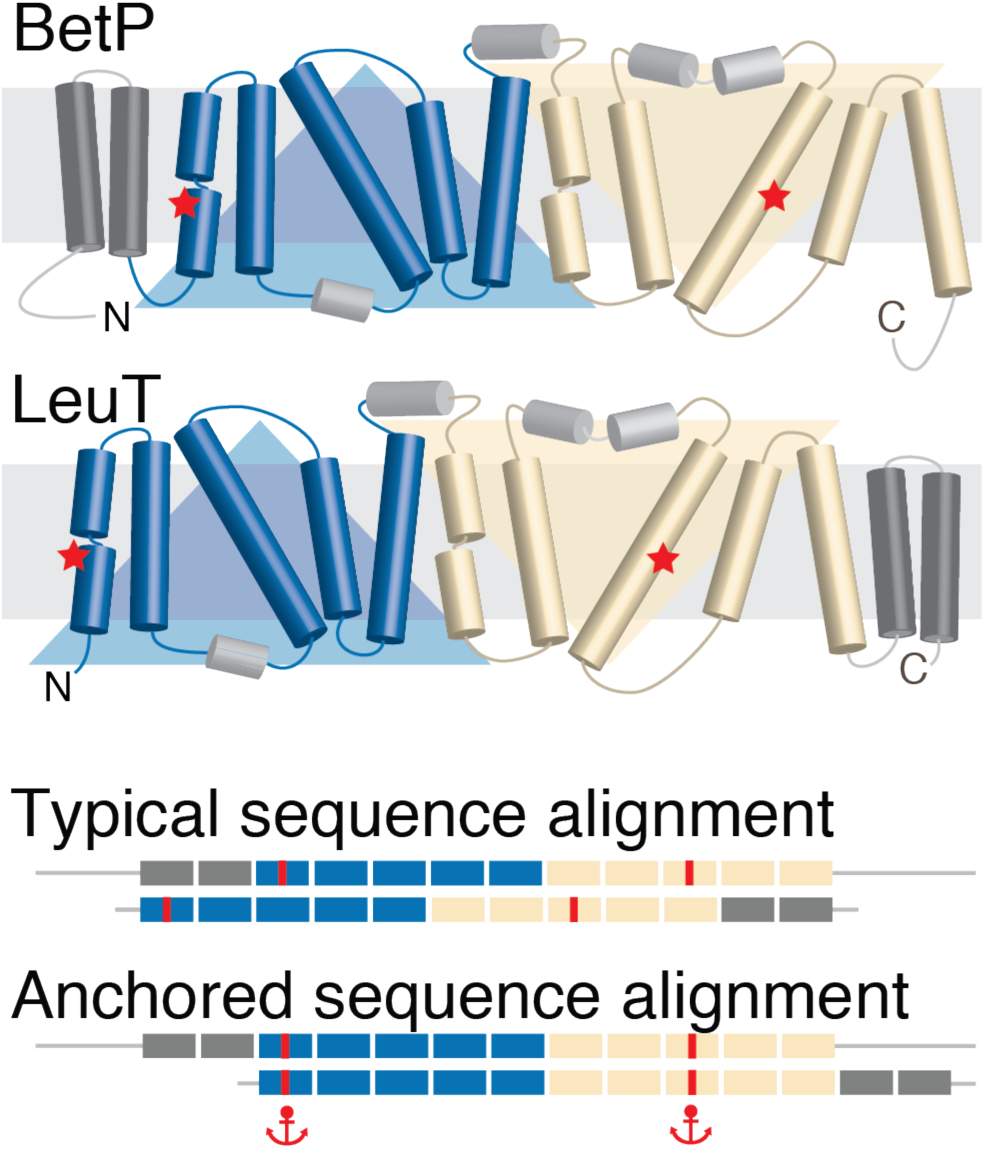
Schematic of the anchors concept as applied to membrane proteins with the so-called LeuT fold. This fold comprises two repeats of five membrane-spanning segments each (*blue and wheat*). The protein BetP contains an additional two transmembrane segments (*dark gray*) N-terminal to the repeats, whereas LeuT contains two additional transmembrane segments at its C-terminal end (*dark gray*). A typical sequence alignment favors matching of all twelve transmembrane segments and thus incorrectly aligns the core segments, including the position of the sodium binding sites (*red*). These errors can be corrected by using anchors (*red*) at specific locations.

Fortunately, in real-life mechanistic studies, it is frequently the case that biochemical or biophysical data is available that identifies a relationship between specific positions in the sequences being aligned. Examples might include the location of binding sites, or of residues key to the folding of the structure. However, unless these residues belong to a larger motif, their signal may be lost among the dissimilarities, and as a result, such positions may not be matched, rendering the final alignment meaningless. A typical solution is to make ad hoc modifications to the alignment that reflect this information. The need for such adjustments is common, for example, in studies where an alignment is required for building a homology model, as correct sequence matching is essential for obtaining an accurate model (Forrest, Tang, and Honig 2006; Luccio and Koehl 2011). However, such interventions reduce the robustness and reproducibility of the remainder of the alignment, as the boundaries of the adjusted regions are determined arbitrarily.

In the absence of a 100% accurate alignment algorithm, an alternative strategy lies in the use of restraints, also referred to as anchors. Some multiple-sequence alignment methods such as DIALIGN (Morgenstern et al. 2005, 2006) and COBALT (Papadopoulos and Agarwala 2007) use information from BLAST or CCD database searches to restrain fragments or domains during the process of constructing full-length protein or genome alignments. Other methods, such as MA-PRALINE (Dijkstra et al. 2018), ConBind (Lelieveld et al. 2016), and FMALIGN (Chakrabarti et al. 2004) allow for matching of regular expressions or sequence motifs. However, requiring the use of sequence motifs fails in cases where only a single residue is conserved, or the signal is weak. The pair-wise profile alignment tool SALIGN (Marti-Renom 2004) in the Modeller suite accounts for this scenario by anchoring positions rather than sequence elements. However, none of the aforementioned methods is designed specifically for high-accuracy pairwise alignments of membrane proteins.

Here, we describe the introduction of restraints as a user-defined option into AlignMe, a method developed specifically for pairwise global alignment of membrane protein sequences (Stamm et al. 2014, 2013). The specificity for membrane proteins was incorporated into AlignMe at two levels. First, optimal forms of input, including sequence homologs, hydrophobicity profiles, secondary structure, and/or transmembrane predictions, were identified based on a data set of membrane proteins of known structure. It was found that the optimal combination of input information differed depending on the similarity of the two proteins being compared: the inclusion of transmembrane predictions, for example, was found to be most impactful for very distantly-related proteins (with <10% sequence identity). Second, the choice of gap penalties was optimized independently for each combination of inputs. Consequently, in practice, AlignMe is a suite of alignment “modes” tailored to the nature of the input sequences (Stamm et al. 2013, 2014).

We test the use of anchors in AlignMe with three case studies similar to those that might be found in functional, modeling, or bioinformatic studies. Our results demonstrate, not only that the anchored positions are correctly matched in the resultant alignment, but also that a single constraint can facilitate a systematic correction across an entire alignment, resulting in dramatic increases in alignment accuracy. Finally, we also illustrate the use of anchors to robustly extend partial alignments of membrane proteins, which, in the case of divergent folds, enables leverage of information from structure-based alignments to obtain more reliable global alignments.

## Materials and Methods

### Anchoring

The Needleman-Wunsch global pairwise alignment algorithm (Needleman and Wunsch 1970) requires the generation of a dynamic programming matrix that reflects the scores of matching every pair of residues in the two sequences based on a scoring function of pre-defined nature. For two sequences of lengths *N* and *M*, this matrix has dimensions (*N*+1) x (*M*+1). The optimal alignment is then selected as the highest-scoring path obtained by tracing back through that matrix.

The ability to anchor positions in the alignment was implemented in AlignMe as follows. First, the dynamic programming matrix is initialized by placing non-zero values, of weight *W*, at the row and column that we wish to anchor, while setting the remainder of the cells to zero. The standard dynamic programming matrix is then constructed on top of this anchored matrix, so that the scores at the anchored positions are increased according to the weights. As the traceback procedure follows the path with the highest scores, it will be biased to pass through the anchored positions. In practice, we impose anchors such that all traceback pathways will pass through this position. Note, however, that in principle, weaker constraints can be used, for example, if there is uncertainty as to the robustness of the relationship between the two positions, or if conflicting information is available.

### SERCA regulatory domain test case

Eight different regulatory subunits of the calcium pump, including *Drosophila sarcolamban* isoform a, dSCLa (UNIPROT codes: C0HJH4, SLCA_DROME) and isoform b, dSCLb (M9PCQ8, M9PCQ8_DROME); human Dwarf Open Reading Frame, hDWORF (P0DN84, DWORF_HUMAN); human myoregulin, hMLN (P0DMT0, MLN_HUMAN); human phospholamban, hPLN (Q5R352, Q5R352_HUMAN), human sarcolipin, hSLN (O00631, SARCO_HUMAN); mouse another regulin, mALN (Q99M08, CD003_MOUSE); and mouse endoregulin, mELN (Q3U0I6, Q3U0I6_MOUSE), were aligned pairwise, using AlignMe in three different modes with default parameters for each mode (Stamm et al. 2013, 2014): (1) Fast mode, which considers hydrophobicity and the VTML substitution matrix (Müller and Vingron 2000); (2) AlignMe P, which considers a position-specific substitution matrix obtained from PSI-BLAST (Altschul et al. 1997); and (3) AlignMe PS, which considers the PSSM as well as secondary structure predictions from PSIPRED (Jones 1999). Note that the fourth available mode, AlignMe PST was not used for this test case. That is because, the PST mode relies on transmembrane predictions from OCTOPUS (Viklund and Elofsson 2008), which did not consistently identify the transmembrane segment in all eight proteins and, as such, not all pairs of alignments could be computed.

Anchors of *W* = 1000 were imposed in each pairwise alignment at four positions in the transmembrane segments shown in **Fig. 2**. The “correct” reference alignment was manually-constructed and reported previously (Primeau et al. 2018). Test alignments were scored relative to the transmembrane domains in this reference alignment, where the transmembrane region is defined as the segment containing residues 31-49 in hPLN. The shift error was measured as the number of positions by which the test alignment was shifted relative to the reference alignment.

**Figure 2.**
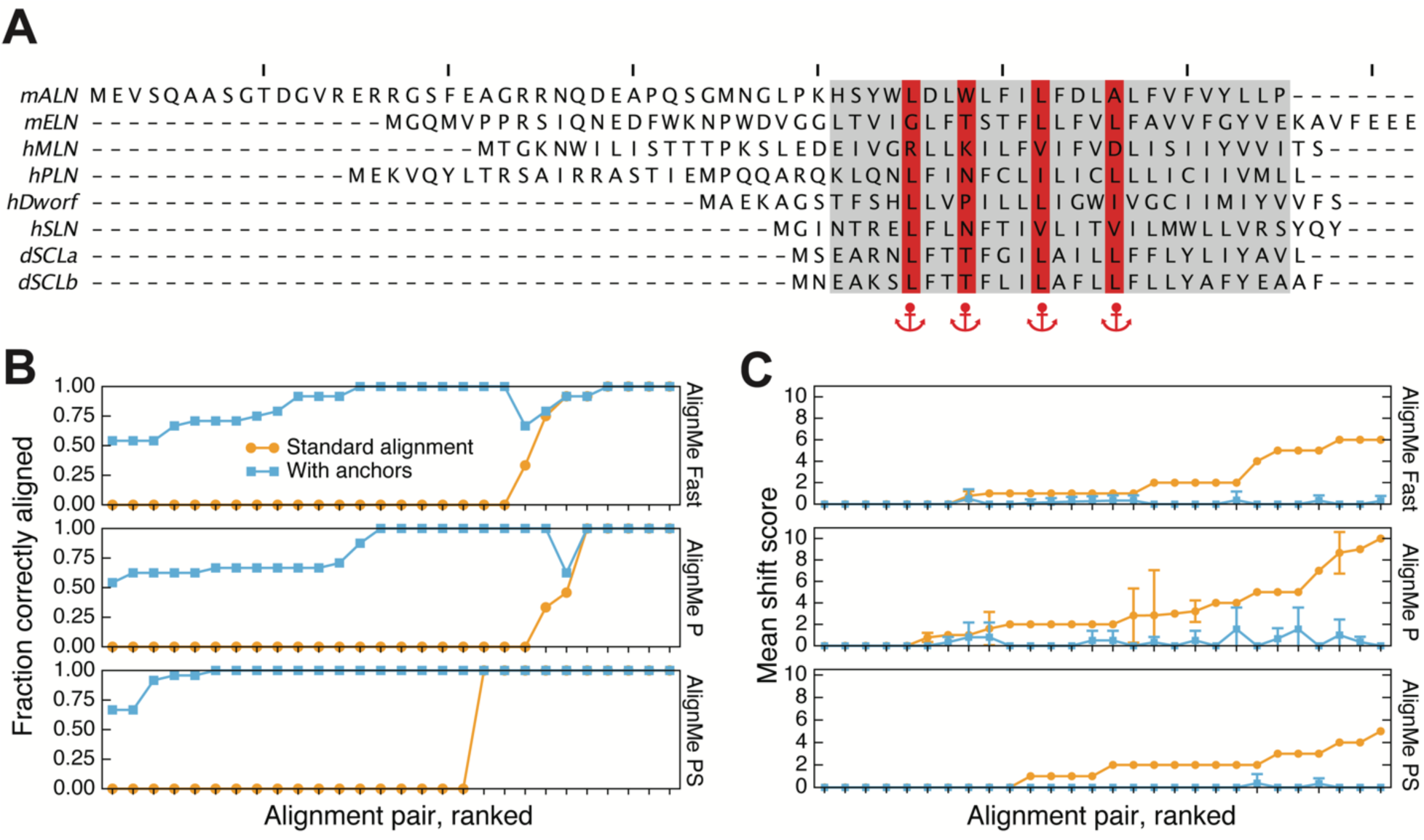
Effect of anchors on the pairwise alignment of SERCA regulatory subunits using AlignMe. **(A)** Multiple sequence alignment of eight SERCA regulatory subunits, namely human phospholamban (hPLN), sarcolipin (hSLN), mouse another-regulin (mALN), mouse endoregulin (mELN), human myoregulin (hMLN), human DWORF (hDWORF), and *Drosophila* sarcolamban a and b isoforms (dSCLa & dSCLb), obtained from (Primeau et al. 2018). Functionally important residues (*red*) and membrane-spanning segment definitions (*gray*) are highlighted, and tick marks indicate every 10 positions along the alignment. **(B, C)** Accuracy of the pairwise alignments obtained either without (*orange circles*) or with (*cyan squares*) anchor constraints imposed at four positions in the transmembrane segment (*red in A*). AlignMe was run using three different modes: fast mode, which considers hydrophobicity and a standard substitution matrix; AlignMe P, which considers a position-specific substitution matrix obtained from PSI-BLAST; and AlignMe PS, which considers the PSSM as well as secondary structure predictions from PSIPRED. **(B)** Fraction of correctly-aligned transmembrane positions for each alignment. Pairs of sequences are sorted according to their accuracy obtained without anchoring. **(C)** The mean number of positions in the transmembrane segments that each aligned column was shifted by, relative to its correct position. Error bars indicate standard deviation.

### LeuT fold test case

Alignments between *Corynebacterium glutamicum* BetP (UniProt entry L7V4D0) and *Aquifex aeolicus* LeuT (UniProt entry O67854) were obtained using AlignMe in PST mode. Residues 150 and 467 in BetP were anchored to residues 23 and 354 in LeuT, respectively. These positions are known to be involved in sodium coordination in the first and eighth transmembrane segments of the core LeuT fold (Yamashita et al. 2005; Khafizov et al. 2012). The reference alignment was a structure-based alignment of the two proteins obtained using SKA (Yang and Honig 2000) on PDB entries 2A65 and 4C7R, for LeuT and BetP, respectively, in addition to AdiC, ApcT, CaiT, Mhp1 and SGLT, in which the core of the fold is superposed (Khafizov et al. 2012). Similar to the SERCA domains, test alignments were scored as the fraction of correctly-aligned residues relative to the reference alignment, excluding any positions in the reference alignment that contain gaps. The shift error was measured as the number of positions by which the test alignment was shifted relative to the reference alignment, either for each column in the alignment or averaged over the length of the alignment.

### LeuT repeats test case

Alignments were carried out between the repeated elements of LeuT, namely TM1-5 (residues 10-234) and TM6-10 (residues 242-443). The reference alignment was a structure-based alignment generated by the symmetry detection method, CE-Symm v2.0 (Myers-Turnbull et al. 2014; Bliven et al. 2019), and anchors were identified in the secondary structure elements from that alignment. Test alignments included increasing numbers of anchors by progressive inclusion of 2-residue windows within secondary structure elements along the sequence, defined as follows: TM1a: 10-24; TM1b: 26-38; TM2: 40-71; intracellular loop 1 (IL1): 76-85; TM3: 87-125; extracellular loop 2 (EL2): 136-153; TM4: 165-184; TM5: 190-214; EL3: 223-232. AlignMe was run in Fast mode (Stamm et al. 2014), which would be a reasonable choice for bioinformatic studies requiring large numbers of alignments. The fraction of correctly-aligned residues was computed only for residues in the transmembrane segments or in secondary-structure elements, as above.

### Software and Availability

The anchoring feature was implemented in AlignMe v1.2, and the code is available through github (https://github.com/Lucy-Forrest-Lab/AlignMe), and as a web service at http://www.bioinfo.mpg.de/AlignMe/.

## Results

### Application of anchors to alignments of SERCA pump regulatory domains

Single-pass, or bitopic, membrane proteins can pose a significant challenge to a sequence alignment method, as their primary feature is a hydrophobic segment, around 25 amino acids in length. Nevertheless, even such simple proteins partake in specific functional interactions, as illustrated by the case of the proteins involved in regulating the **S**arco/**E**ndoplasmic **R**eticulum **C**alcium **A**TPase, also known as **SERCA**, the calcium pump, or Ca^2+^-ATPase. These regulatory subunits, which include phospholamban and sarcolipin, modify the functional properties of SERCA through direct interactions involving their transmembrane subunits.

To illustrate the challenge posed by these sequences, pairwise alignments of eight known SERCA regulatory domains were obtained using AlignMe in several different modes (Fast, P and PS), but without anchoring. Accuracy was measured as the fraction of aligned positions that agree with those in the transmembrane region of the previously-reported alignments, for each pair of sequences (**Fig. 2A**). Although the accuracy depended slightly on the alignment mode used, only around a third of the pairwise alignments were accurate for >50% of their positions (**Fig. 2B**), while the remainder were misaligned along the full length of the transmembrane regions.

Since the metric of absolute accuracy does not provide information about the extent by which the alignment is wrong, we also measured the mean shift score, i.e., the number of positions by which a matched pair is offset from its expected position, averaged along the length of the sequence. When no anchors were included in the alignments, the mean shift errors were consistently around 3, i.e. a turn of a helix, but were also found to be as high as 10 (**Fig. 2C**).

Previous studies have demonstrated that four residues in the transmembrane segments of phospholamban and sarcolipin are essential for their regulatory function (for a review, see (Primeau et al. 2018)), and thus contribute to the interface with the SERCA pump. Constraining these four positions in each of the pairwise alignments resulted in a dramatic increase in accuracy: in all 27 pairs, >50% of the positions in the predicted transmembrane segment were aligned correctly (**Fig. 2B**), relative to the manually-constructed pairwise alignments (**Fig. 2A**). Moreover, the extent of the shift error was dramatically reduced using this approach, with mean values of <1.5, suggesting that, even for positions identified as strictly incorrect, AlignMe always placed them within 2 or three positions of that in the reference alignment (**Fig. 2C**). We note that it is also possible that those reference alignments contain minor errors, especially at positions far from the anchor points. Thus, the new, anchored alignments, which rely exclusively on experimental and evolutionary information, could potentially contain mechanistic insights regarding the SERCA regulatory subunits.

### Application of anchors to alignments of LeuT fold transporters

The above examples are limited by a lack of direct structural data. Moreover, the short length of the transmembrane segments in these bitopic proteins proved a challenge to the transmembrane prediction tool OCTOPUS, and thus no alignments could be obtained using AlignMe’s PST mode. We therefore also illustrate the effect of anchors on another system, namely structures belonging to the APC superfamily of transporters, whose structures contain the so-called LeuT fold (see **Fig. 1**). In the case of the proteins LeuT and BetP, high-resolution structures are available that can be used to create a reference structural alignment. Moreover, biochemical and biophysical experiments have demonstrated a common functional property of these proteins, namely, a conserved sodium binding site involving side-chain and backbone groups in the first and eighth transmembrane segments of the core fold.

We first demonstrate that a standard alignment approach, even using the advanced features of AlignMe PST, does not even match the transmembrane segments in the two repeats (**Fig. 3A**), let alone the sodium binding site residues. Thus, the resultant alignment is wildly incorrect for the entire length of the protein, with each pair of residues shifted by >200 from their correct positions (**Fig. 4**, orange). By contrast, imposing constraints at positions in the sodium binding site allows AlignMe to correctly match the transmembrane core segments (**Fig. 3B**,**3C**) such that the majority of the positions in the >450-residue long alignment are within five positions of the locations expected based on the structure alignment (**Fig. 4**).

**Figure 3.**
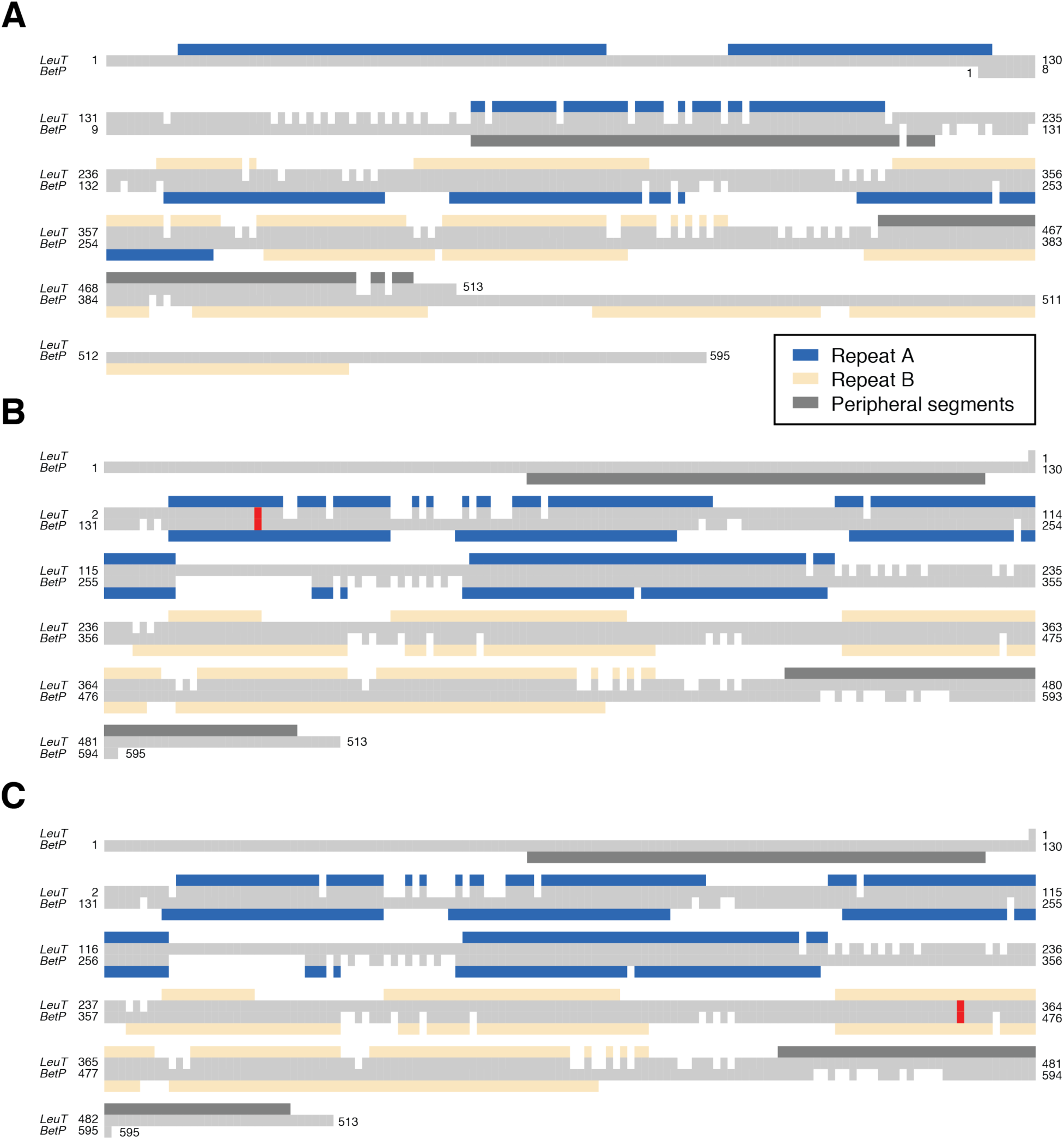
Alignments between two structurally-related membrane proteins, BetP and LeuT. Alignments were obtained either **(A)** with a standard protocol or **(B, C)** with a single anchor imposed (*red*). Anchors were placed between residues involved in coordinating sodium ions in the **(A)** first or **(B)** eighth transmembrane segments of the core LeuT fold. The transmembrane segments of the two proteins are indicated using boxes above and below the respective sequences, highlighting the structurally-related elements, repeat A (*blue*) and repeat B (*wheat*), as well as the transmembrane segments that are peripheral to the core fold (*dark gray*). See Figure 1 for more information.

**Figure 4.**
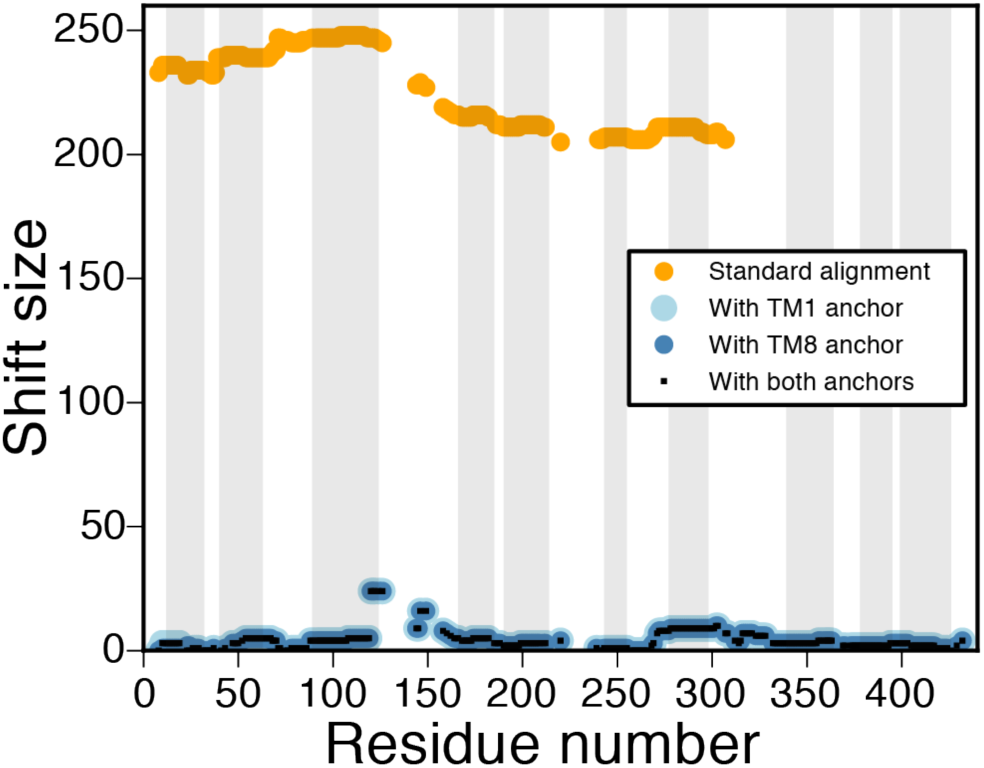
Comparison of the effect of anchors on the alignments of BetP and LeuT. The shift error, relative to the structure-based alignment, is plotted as a function of the residue number in LeuT, for alignments obtained either with the standard methodology (*orange*) or with anchors imposed (*light blue, dark blue or black*). Anchors were placed between residues involved in coordinating sodium ions clustered in the first and eighth transmembrane segments of the core LeuT fold, either separately or simultaneously. The approximate transmembrane regions of LeuT are indicated by gray boxes.

Moreover, these improvements were achieved independent of whether the constraint was located in TM1, in TM8, or in both.

### Assessment of the compounding effects of anchors

The above examples illustrate how just a few anchors can dramatically improve pairwise alignments of distantly related proteins, and therefore could be insightful for mechanistic and evolutionary studies of their function, or beneficial for structural modeling studies. Another promising application of anchors is situations in which a full-length alignment of two sequences is required, but where only fragments of the alignment are available from other tools. Such sources of aligned fragments might include local alignments, or structural alignments of repeated elements within a structure, as in the case of membrane-protein symmetry analysis (Sarti et al. 2018; Aleksandrova, Sarti, and Forrest 2018). In such cases, it is desirable to have an unbiased method by which to construct the remainder of the alignment.

As a test of such a situation, we examined the case of the internal repeats in LeuT, whose sequences contain <10% identical residues (Forrest et al. 2008). Specifically, we aligned the two five-transmembrane-helix repeats (**Fig. 1**) using constraints within the secondary structure elements that were selected based on the known symmetry relationship (**Fig. 5**). The reference alignments used to define the anchors and to assess accuracy of the alignments were obtained using a symmetry-detection algorithm, CE-Symm. Using all the symmetry-based restraints, all the positions in the alignment were correct (**Fig. 5, red**). By contrast, a control alignment, generated using AlignMe without any anchors, only contains 8.1% of its positions correctly aligned (**Fig. 5, black**). We note that, as constraints are progressively removed from the list, starting in the N-terminal segment (TM1a), the accuracy progressively decreases, albeit with a few abrupt transitions (**Fig. 5, colors**). Together with the data for BetP and LeuT, these results suggest that, for very distantly-related sequences, either large numbers of, or judiciously-placed constraints may be important for obtaining an accurate alignment.

**Figure 5.**
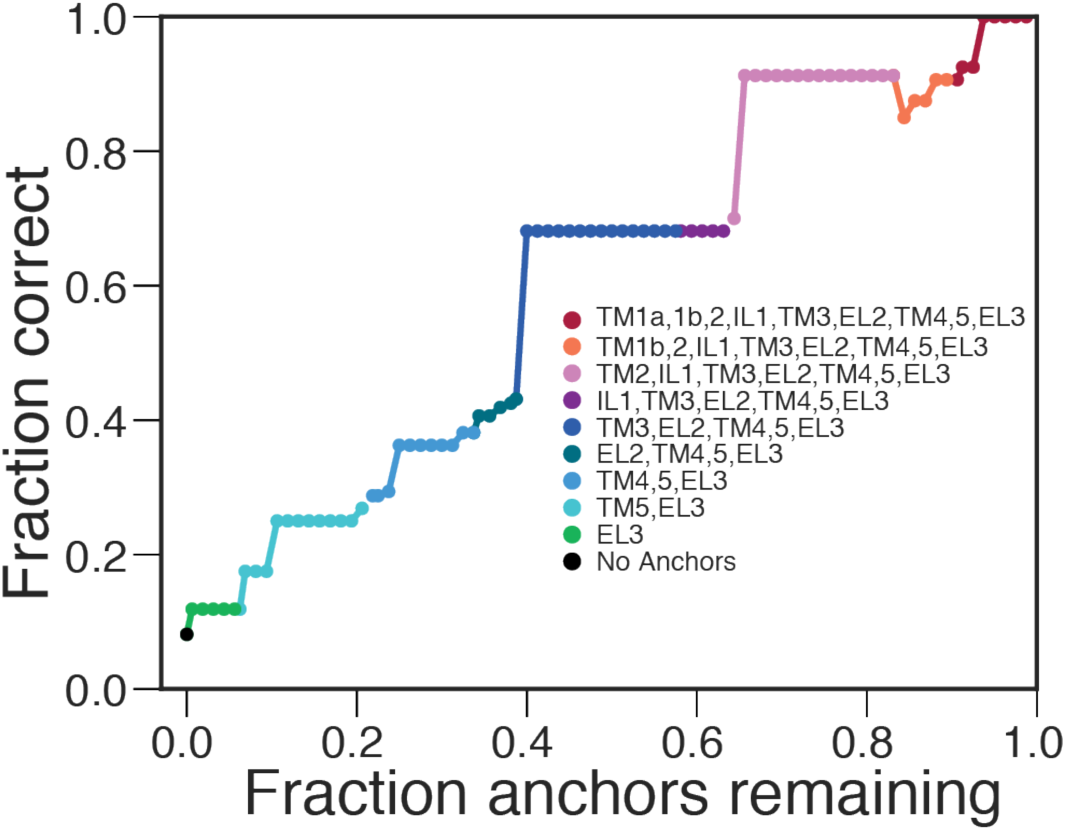
Effect of different numbers of anchors on alignments of the internal repeats of LeuT, as measured by the fraction of the alignment that is correctly aligned. Constraints were imposed within secondary structure elements of the first repeat, TM1a, TM1b, TM2, IL1, TM3, EL2, TM4 TM5 and EL3, to match with their equivalent positions in the second repeat, based on a structure-based alignment obtained using the symmetry-detection algorithm CE-Symm. The effect of adding increasingly large subsets of restraints is shown relative to the accuracy of the alignment obtained without any anchors (*black*).

## Discussion

In this work, we addressed the challenging problem of aligning membrane protein sequences. We implemented the proposed anchoring approach in the AlignMe software package, as its features were designed and optimized specifically with membrane proteins in mind. However, anchors could in principle be applied to the alignment of any pair of primary sequences within the framework of a Needleman-Wunsch algorithm. Indeed, since the framework of the AlignMe software is such that it can read any type of input information or set of gap penalties, AlignMe itself could readily be repurposed using standard position specific substitution matrices or general substitution matrices such as BLOSUM, for anchoring in any pair of sequences.

The example of BetP and LeuT illustrates a situation involving large multi-spanning proteins, where the presence of two peripheral segments in different positions can stymie even the most robust of global alignment methods. This is because global alignment methods favor matching of all transmembrane segments simultaneously. In this case, a single constraint has the power to dramatically improve the alignments.

In addition to functional studies of specific proteins, anchors may have a number of other applications in bioinformatic workflows. For example, they can be used to expand a robust, local alignment, such as that obtained from a structure-based, or motif-matching approach, into a full-length alignment suitable for situations where the entire protein sequence is required (Morgenstern et al. 2006; Papadopoulos and Agarwala 2007). In future work then, we plan to utilize the anchoring strategy in AlignMe in completing global alignments of internal structural repeats obtained by structure-based symmetry detection methods such as CE-Symm, so that those alignments can be readily compared across structures and family members of membrane proteins in the EncoMPASS database (Sarti et al. 2018).

In the applications provided here, the strength of the anchors imposed was deliberately set to a very large number to ensure that they would override all other paths in the dynamic programming matrix, and thereby essentially fix the alignment at that point. Nevertheless, it could be of interest to explore the effect of varying the strengths of the anchors. For example, one might imagine situations in which candidate anchors are mutually exclusive, and therefore the ability to violate one or more anchors would be of value.

In conclusion, anchoring is a powerful addition to the arsenal of sequence alignment approaches, and its implementation within AlignMe makes it readily available to those interested in carrying out alignments of membrane proteins.

## Acknowledgements

This research was supported by the Division of Intramural Research of the NIH, National Institute of Neurological Disorders and Stroke and by the Max Planck Gesellschaft. We thank Drs. Mykola Petrov and Markus Rampp for their support of the MPG supercomputing center and the AlignMe server.

## Author Contributions

Conceptualization: AAA, KK, RS, and LRF; Methodology: RS, ES, and AAA; Development of AlignMe code: RS and KK; Development of AlignMe web server: RS, ES, MS, and LRF; Validation, ES, and LRF; Code development and analysis of test cases: ES, EY, AAA, and LRF; Resources, Project administration and Funding acquisition: LRF; Data curation: ES, EY and LRF; Writing of original draft: RS, KK and LRF; Reviewing and editing drafts: all authors; Visualization of data: ES, EY and LRF; Supervision: AAA, RS, and LRF.

